# Associations between SARS-CoV-2 Infection or COVID-19 Vaccination and Human Milk Composition: A Multi-Omics Approach

**DOI:** 10.1101/2024.04.12.589299

**Authors:** Sneha Couvillion, Ernesto S. Nakayasu, Bobbie-Jo M. Webb-Robertson, Isabella Yang, Josie Eder, Carrie D. Nicora, Lisa M. Bramer, Yuqian Gao, Alisa Fox, Claire DeCarlo, Xiaoqi Yang, Mowei Zhou, Ryan M. Pace, Janet E. Williams, Mark A. McGuire, Michelle K. McGuire, Thomas O. Metz, Rebecca L. Powell

## Abstract

The risk of contracting SARS-CoV-2 via human milk-feeding is virtually non-existent. Adverse effects of COVID-19 vaccination for lactating individuals are not different from the general population, and no evidence has been found that their infants exhibit adverse effects. Yet, there remains substantial hesitation among this population globally regarding the safety of these vaccines. Herein we aimed to determine if compositional changes in milk occur following infection or vaccination, including any evidence of vaccine components. Using an extensive multi-omics approach, we found that compared to unvaccinated individuals SARS-CoV-2 infection was associated with significant compositional differences in 67 proteins, 385 lipids, and 13 metabolites. In contrast, COVID-19 vaccination was not associated with any changes in lipids or metabolites, although it was associated with changes in 13 or fewer proteins. Compositional changes in milk differed by vaccine. Changes following vaccination were greatest after 1-6 hours for the mRNA-based Moderna vaccine (8 changed proteins), 3 days for the mRNA-based Pfizer (4 changed proteins), and adenovirus-based Johnson and Johnson (13 changed proteins) vaccines. Proteins that changed after both natural infection and Johnson and Johnson vaccine were associated mainly with systemic inflammatory responses. In addition, no vaccine components were detected in any milk sample. Together, our data provide evidence of only minimal changes in milk composition due to COVID-19 vaccination, with much greater changes after natural SARS-CoV-2 infection.

**IMPORTANCE:** The impact of the observed changes in global milk composition on infant health remain unknown. These findings emphasize the importance of vaccinating the lactating population against COVID-19, as compositional changes in milk were found to be far less evident after vaccination compared to SARS-CoV-2 infection. Importantly, vaccine components were not detected in milk after vaccination.

## BACKGROUND

Human milk is considered the gold standard for infant nutrition, and all leading health organizations recommend human milk-feeding exclusively for 6 months and alongside other forms of nutrition for at least 2 years [1]. Human milk provides individualized combinations and concentrations of all the essential nutrients as well as a host of immune cells, microbes, and immunomodulatory components that provide protection to the infant during a period characterized by an immature immune system. As such, there is substantial evidence demonstrating the benefits of human milk in protecting infants against a wide range of infections [2, 3]. Despite its benefits, human milk can also serve as a vehicle of vertical transmission for a small subset of pathogens, including HIV, Zika, and Ebola viruses [4, 5]. At the onset of the COVID-19 pandemic, there was a major concern regarding potential SARS-CoV-2 transmission via human milk-feeding. This resulted in various public health organizations recommending that SARS-CoV-2-infected people avoid chest/breastfeeding and/or separate from their infants, often with adverse consequences to establishment of the chest/breastfeeding relationship, in some cases irrevocably [6]. In response, and to characterize the milk immune response to SARS-CoV-2 infection, we and others analyzed milk obtained during and after SARS-CoV-2 infection for viral RNA, viable SARS-CoV-2, and SARS-CoV-2-specific antibodies [7–9]. Although a handful of studies identified SARS-CoV-2 RNA in a small number of milk samples, the preponderance of evidence indicated that milk is not a vehicle of SARS-CoV-2 transmission [10–12].

Amid determining whether human milk-feeding was safe while the lactating parent was infected with SARS-CoV-2, researchers and the public simultaneously faced a second global crisis related to human milk-feeding and the COVID-19 pandemic: the novel COVID-19 vaccines were not evaluated for safety in the lactating population. As such, some countries recommended that these vaccines not be offered to lactating people, and there was and remains hesitation among this population globally regarding the safety of these vaccines for their infants [13, 14], even with the finding that particularly the mRNA-based vaccines induced a robust antibody response in human milk [15, 16] .

To date there have been limited reports comparing differences in milk composition associated with SARS-CoV-2 infection, and importantly there remains a substantial knowledge gap concerning the impact of COVID-19 vaccines on human milk composition. To our knowledge, there are only two peer-reviewed reports of detailed examinations of the molecular composition of milk following infection, and no published studies on compositional changes due to vaccination. Zhao *et al.* reported alterations in the milk proteome and metabolome and Arias-Borrego and co-authors examined changes in minerals and metabolites, all after infection with SARS-CoV-2 [17, 18]. Here we employed a more holistic multi-omics (metabolomic, proteomic, and lipidomic) approach to: 1) characterize and compare the composition of milk produced by SARS-CoV-2-infected participants to that produced by participants prior to vaccination; and 2) characterize and compare composition of milk produced before and after the first dose of Pfizer, Moderna, Johnson and Johnson (Janssen; J&J) COVID-19 vaccines, using each vaccinated person as their own control.

## RESULTS

Study participants were 26 to 41 years old (average of 32 years) and from <1 to 30 months postpartum (average of 8 months) (Table S1). Analytes were extracted from milk samples using MPLEx [19] and analyzed using mass spectrometry-based proteomics, metabolomics, and lipidomics (see Methods for details). Tables S2 - S4 list detailed data on compounds exhibiting significant changes compared to control samples.

### Association of SARS-CoV-2 Infection and Changes in Milk Composition

#### Proteomics

SARS-CoV-2 infection was associated with changes to 67 proteins in milk compared to control samples in the 7 days following infection, with 43 proteins exhibiting significant increases and 24 proteins exhibiting significant decreases (Fig. 1). Pathway enrichment analysis indicated that these proteins were mainly associated with systemic inflammatory responses to SARS-CoV-2 and various other infections (e.g. hepatitis C, measles, influenza A) and inflammatory conditions (e.g. NOD-like receptor signaling, JAK-STAT signaling and inflammatory bowel disease) (Fig. 1B). The last regulated pathway was the aldosterone synthesis and secretion pathway (Fig. 1).

**Figure 1:**
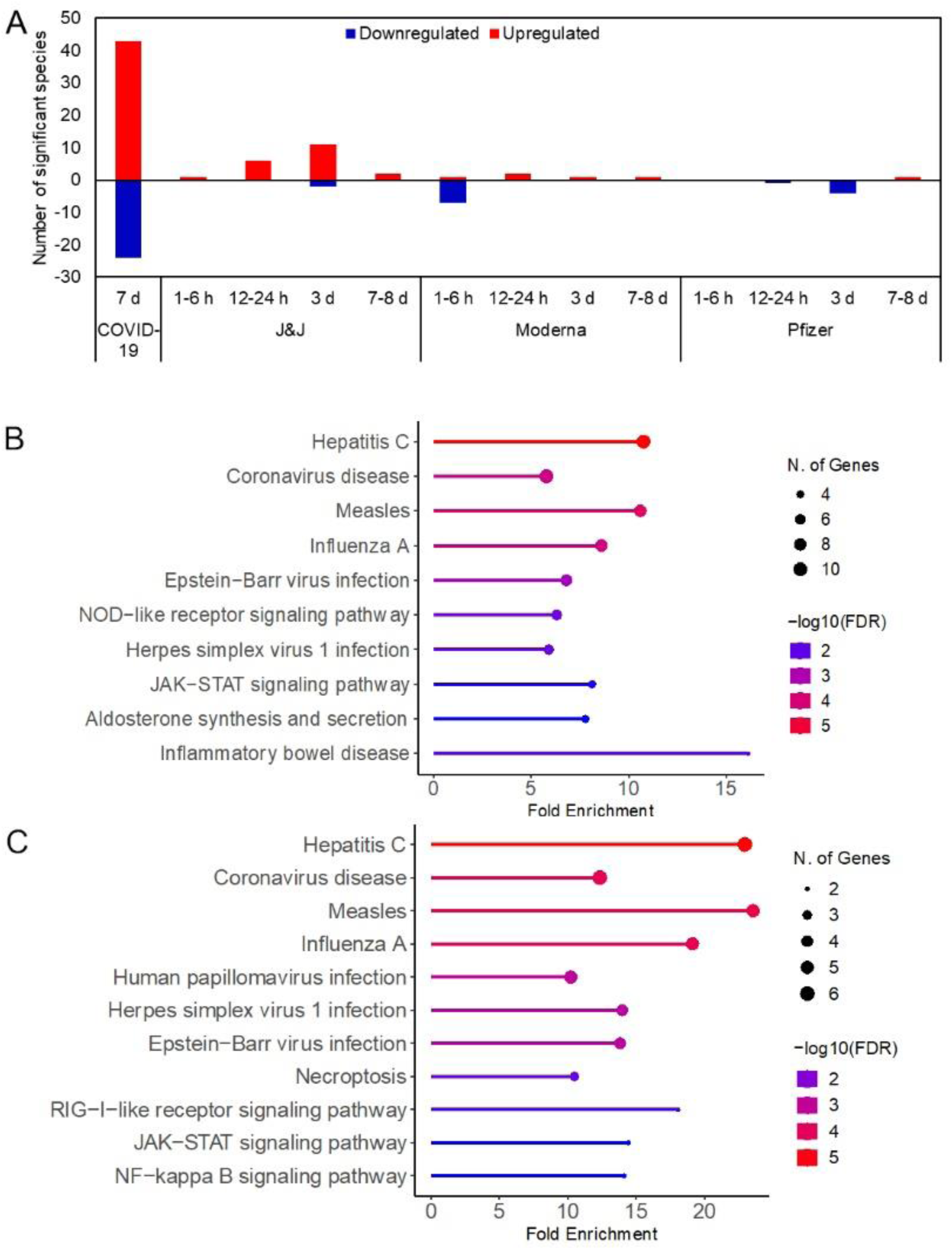
COVID-19 vaccination results in minimal impact on milk proteomic composition in comparison to SARS-CoV-2 infection. (A) Number of proteins that significantly increased (red) or decreased (blue) in milk produced after SARS-CoV-2 infection (COVID-19) or COVID-19 vaccination. All comparisons were made to the pre-dose time point. (B-C) Pathway enrichment analysis indicates altered milk proteins are related to systemic inflammatory and immunomodulatory pathways. (A) SARS-CoV-2 infection. (B) Post J&J vaccination. The pathway enrichment analysis included significant proteins from all 4 time points.

#### Lipidomics

SARS-CoV-2 infection was associated with changes to 385 lipid molecular species in milk (195 increased and 190 decreased) (Fig. 2). These lipids were from 5 classes and 21 subclasses of lipids (Fig. 2C). Of note, 66 species of triacylglycerols increased, while 61 fatty acids and 57 diacylglycerols decreased (Fig. 2). Among the glycerophospholipids, phosphatidylethanolamine and lysophosphatidylethanolamine were the most regulated subclasses with 44 and 19 upregulated lipids, respectively (Fig. 2). There was also a reduction in 20 fatty acid esters of hydroxyl fatty acids (generally anti-inflammatory lipids) and an increase in 8 ceremide species (generally considered pro-inflammatory lipids) (Fig. 2). These results show that the SARS-CoV-2 infection likely impacts milk lipid composition, in general elevating pro-inflammatory lipids and reducing anti-inflammatory ones.

**Figure 2:**
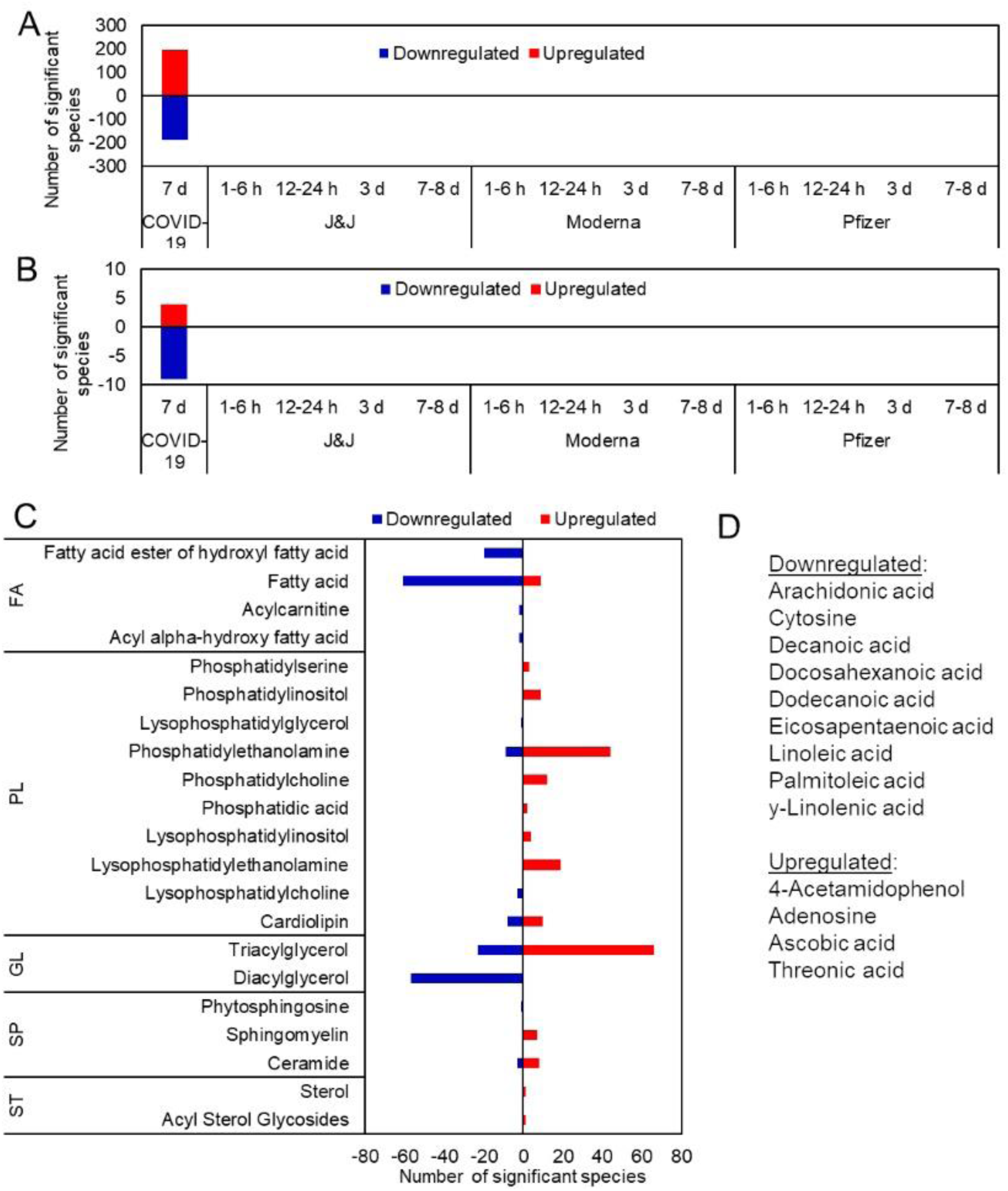
COVID-19 vaccination results in minimal impact on milk lipid and metabolic composition in comparison to SARS-CoV-2 infection. Number of (A) lipids and (B) metabolites that significantly increased (red) or decreased (blue) after SARS-CoV-2 infection (COVID-19) or COVID-19 vaccination. (C) Lipid classes that were regulated in SARS-CoV-2 infection. (D) List of metabolites that were regulated in SARS-CoV-2 infection. FA: fatty acid; PL: (glycero)phospholipid; GL: glycerolipids; SP: sphingolipid; ST: sterol.

#### Metabolomics

SARS-CoV-2 infection was associated with changes to 13 metabolites in milk (4 increased, 9 decreased) (Fig. 2). Eight of the 9 downregulated metabolites were fatty acids (Fig. 2), cross-validating the lipidomics observations described above. Of note, 4-acetamidophenol (also known as the anti-inflammatory drug acetaminophen), and ascorbic acid (vitamin C) and its degradation intermediate, threonic acid, were upregulated after infection (Fig. 2).

### Association of COVID-19 Vaccination and Changes in Milk Composition

#### Proteomics, Lipidomics and Metabolomics

Vaccination was not associated with any changes in lipid or metabolite evaluated (Fig. 2). However, milk proteomic profiles differed by vaccination type and timing (Fig. 1). At 1-6 h, milk produced by women receiving the Moderna vaccine exhibited changes in 8 proteins, while those receiving the J&J vaccine exhibited a change in only 1 protein (Fig. 1). Milk composition did not change after receiving the Pfizer vaccine (Fig. 1). At 12-24 h, J&J exhibited changes to 6 proteins, while Moderna and Pfizer exhibited changes to 2 and 1 proteins, respectively (Fig. 1). At day 3, milk produced by participants receiving the J&J vaccine exhibited changes to 13 proteins, while that produced by participants receiving the Moderna and Pfizer vaccines exhibited changes to 2 and 4 proteins, respectively (Fig. 1). At day 7, milk produced by participants receiving the J&J vaccine exhibited changes to 2 proteins, while that produced by participants receiving the Moderna and Pfizer vaccines exhibited changes to 1 protein each (Fig. 1).

Only the J&J vaccine resulted in changes sufficient to perform a pathway enrichment analysis. Similar to SARS-CoV-2 infection, enriched proteins were associated with systemic inflammatory responses to SARS-CoV-2 and various other infections and inflammatory conditions (Fig. 1). Notably, unlike infection, vaccination was not associated with the NOD-like receptor signaling, aldosterone synthesis and secretion, and inflammatory bowel disease pathways (Fig. 1). Conversely, vaccination was also associated with the herpesvirus, necroptosis, RIG-I-like receptor signaling, and NF-kappa B signaling pathways (Fig. 1).

### Evaluation of milk samples for the presence of vaccine components

To assess if vaccine components enter the milk, we investigated the presence of unique components of each vaccine in the samples. For the mRNA vaccines, we searched for the synthetic lipid compound ALC-0315 in the Pfizer vaccine and SM-102 in the Moderna vaccine (Sup Figs. 1-2). For the J&J vaccine, we performed de novo sequencing of the peptides in the J&J vaccine itself to determine viral protein sequences. No adenoviral proteins were detected in any of the milk samples. To conclude, despite the mass spectrometry sensitivity no vaccine components were detected in the milk samples.

## Discussion

There is now global consensus that the risk of contracting COVID-19 via human milk-feeding is virtually non-existent, and the benefits of continuing to chest/breastfeed during and after infection and vaccination are substantial [20, 21]. Adverse effects of vaccination for lactating individuals are generally mild and transitory and not different from the general population, and there exists no clear evidence that infants fed milk produced by recently-vaccinated people exhibit adverse effects [22, 23]. While several publications have reported the presence of trace amounts of vaccine mRNA in a small subset of milk samples [24, 25], the physiological significance remains undetermined. Nonetheless, there remains a substantial hesitancy among some lactating individuals regarding COVID-19 vaccination.

In the present study we used advanced ‘multi-omics’ approaches to determine if compositional changes (including vaccine components) in milk occur after SARS-CoV-2 infection or vaccination. Our extensive analyses showed that multiple compositional changes occur in milk produced within 7 days of confirmed SARS-CoV-2 infection. These changes are highly consistent with a systemic inflammatory response associated with SARS-CoV-2 and various other infections. Interestingly, we also identified both acetaminophen and vitamin C in milk produced by SARS-CoV-2 infected individuals, which likely reflects consumption by participants to alleviate symptoms of illness. To the best of our knowledge, this is the first study to demonstrate such changes occurring in human milk. While SARS-CoV-2 infection elicited changes to over 65 milk proteins, vaccines elicited changes in 13 or fewer proteins at any time point. Notably, the kinetics of the changes measured in milk produced following vaccination were minimal, with Moderna elicited changes in only 8 proteins at 1-6 h after vaccination, while Pfizer changes in 4 proteins at 3 days after vaccination, their highest points. The J&J adenovirus-vectored vaccine was designed to elicit an anti-viral inflammatory response as adjuvant to and vehicle of SARS-CoV-2 immunogen. Consequently, it was not surprising that, compared to the mRNA vaccines, the J&J vaccine elicited the most change in milk composition, (13 changed proteins at 3 days). SARS-CoV-2 infection induced changes in 395 lipids and 13 metabolites, while none of the vaccines induced changes in the lipidomic and metabolomic profiles. The impact on infant health of these changes remains unknown; however, these findings emphasize the importance of vaccinating lactating individuals against COVID-19, as compositional changes in milk were far fewer than those observed after vaccination. Furthermore, we found no evidence of vaccine components in any milk sample produced after vaccination.

## Acknowledgments

Funding for this project was obtained from the National Science Foundation (IOS-BIO 2031753 and 2031715), the Bill and Melinda Gates Foundation (INV-016943), and the Icahn School of Medicine at Mount Sinai. A portion of the research was performed on a project award (doi.org/10.46936/ltds.proj.2021.60066/60000403) from the Environmental Molecular Sciences Laboratory (EMSL), a U.S. Department of Energy Office of Science User Facility located at Pacific Northwest National Laboratory (PNNL) and sponsored by the Biological and Environmental Research program. COVID-19 vaccines were generously donated by the Idaho Department of Health and Welfare. We are grateful to all the study participants and the following individuals who assisted in sample collection: Dr. Kimberly Lackey, Christina Pace, Beatrice Caffé, and Alexandra Navarrete. Omics analyses were performed in the EMSL. Battelle operates PNNL for the DOE under contract DE-AC05-76RLO01830.

## MATERIALS AND METHODS

### Participant Information and Milk Collection Procedures

Participants in this study have been previously described [8, 16]. All participants gave informed consent. Participants receiving a vaccine were eligible to have their milk samples included in this analysis if they were ≥18 years of age, lactating, had no history of a suspected or confirmed SARS-CoV-2 infection, and were scheduled to be vaccinated with the Pfizer, Moderna, or J&J COVID-19 vaccines (original formulations available in 2021). Participants were asked to collect ∼30mL of milk per sample into a clean container according to the requested schedule using electronic or manual pumps in their homes. Samples were collected before the vaccination and 1-6 h, 12-24 h, 3 d and 7 d after the vaccination. Sample collection procedures for these individuals was approved by the institutional review board (IRB) at Mount Sinai Hospital (IRB 19-01243).

Infection study participants were eligible to have their milk samples included in this analysis if they were ≥18 years of age, lactating, and had tested positive for SARS-CoV-2 infection in the previous 7 days. Sample collection procedures for this cohort was approved by the Institutional Review Board (IRB) at the University of Idaho (IRB 20-056 & 20-060). Participants self-collected up to 30 mL of milk using provided sterile, single use collection kits. All milk was frozen in participants’ home freezers until transferred via local transportation or shipping; milk was then stored at -80 °C.

### Protein, Metabolite and Lipid Extraction (MPLEx)

Aliquots of NIST SRM 1953 Organic Contaminants in Non-fortified Human Milk (NIST, Gaithersburg, MD) were used as quality control (QC) samples. Milk samples were blocked and randomized into sample batches of 30 along with a QC milk aliquot. Milk batches were allowed to thaw for ∼45 min inside a biological safety cabinet (BSC) and extracted using MPLEx according to standard protocol [19]. Notably, MPLEx inactivates pathogenic viruses and bacteria with exposed lipid layers, including Middle Eastern Respiratory Syndrome coronavirus [26]. A 1 mL cold (-20 °C) cholorform:methanol working mix (prepared 2:1 (v/v) along with lipid-internal standard (IS) TG 60:1 (20:0-20:1-20:0 D5-TG); TG 44:1 (14:0-16:1-14:0 D5-TG) (Avanti Polar

Lipids, Birmingham, AL) and metabolite-IS 13C-glucose, 13C-sorbitol, 13C-succinic acid (Sigma, St. Louis, MO) was pipetted into a chloroform compatible 2-mL Sorenson MulTI™ SafeSeal™ microcentrifuge tubes (Sorenson bioscience, Salt Lake City, UT) inside an ice-block. Cold water and 50 µL of milk sample was added to make a final ratio of 8:4:3 chloroform:methanol:water/sample, vortexed and allowed to incubate in the ice block for 15 min. The samples were centrifuged at 12,000 *xg* for 10 min to separate the polar, non-polar phases and the protein interlayer. Metabolite samples included 400 µL of the upper polar phase and 400 µL of the lower non-polar phase. The lipid samples included 400 µL of the lower non-polar phase.

The remaining protein interlayer was rinsed with 500 µL of cold 100% methanol, vortexed and centrifuged for 5 mins to pellet the protein and the wash layer was added to the metabolites vial. Protein pellets were lightly dried under a nitrogen stream and stored at -80°C. Metabolite and lipid samples were dried in a vacuum centrifuge and 500 µL 2:1 (v:v) cold chloroform:methanol was added to the lipids; this was stored along with the dry metabolites at -20°C.

### Protein Digestion

The protein interlayer was processed by adding 200 µL of 8M urea in 100 mM triethylamonium bicarbonate, pH 8.0 (TEAB) to the protein pellets and vortexed into solution. A bicinchoninic acid assay (Thermo Scientific, Waltham, MA USA) was performed to determine protein concentration for quality control. Following the assay, 10 mM dithiothreitol was added to the samples and incubated at 60 °C for 30 min with constant shaking at 800 rpm. Reduced cysteine residues were alkylated by adding 400 mM iodoacetamide (Sigma-Aldrich) to a final concentration of 40 mM and incubating in the dark at room temperature for 1 h. Samples were then diluted 8-fold to prepare for digestion with 100 mM TEAB, 1 mM CaCl_2_ and sequencing-grade modified porcine trypsin (Promega, Madison, WI) was added to all protein samples at a 1:50 (w/w) trypsin-to-protein ratio for 3 h at 37 °C. Digested samples were desalted using a 4-probe positive pressure Gilson GX-274 ASPEC™ system (Gilson Inc., Middleton, WI) with Discovery C18 100 mg - 1 mL solid phase extraction tubes (Supelco, St. Louis, MO), using the following protocol: 3 mL of methanol was added for conditioning followed by 2 mL of 0.1% trifluoroacetic acid (TFA) in water. The samples were acidified to 0.1% TFA then loaded onto each column followed by 4 mL of 95:5 water:acetonitrile, 0.1% TFA . Samples were eluted with 1 mL 20:80 water:acetonitrile, 0.1% TFA. The samples were concentrated to ∼100 µL using a vacuum centrifuge.

### Tandem Mass Tag (TMT) Isobaric labeling

A bicinchoninic acid assay was performed to determine the peptide concentration for TMT Isobaric labeling. To generate a common universal reference, 2 µg of peptide from each sample was combined and the pool was assayed for accuracy. Note that in large-scale studies that compare samples across multiple TMT sets, distribution plots can be generated by calculating the peptide intensity ratios of each channel to a common universal reference within each set [27].

For each TMT set, 30 µg of the pooled reference and 30 µg from each sample was aliquoted into new tubes for labeling and dried completely in a vacuum centrifuge.

Samples were randomized into 20 sets including a global pool in each plex to ensure correlation across sets. Each sample was diluted in 40 µL 500 mM HEPES, pH 8.5 [28] and were labeled using amine-reactive 16-plex TMT kits (Thermo Scientific, Rockford, IL) according to the manufacturer’s instructions. Briefly, 250 μL of anhydrous acetonitrile was added to each 5 mg reagent, vortexed and allowed to dissolve for 5 min with occasional vortexing. Reagents (10 µL) were then added to each sample and incubated for 1 h at room temperature with shaking at 400 rpm. Each sample was then diluted with 30 µL 20% acetonitrile. A portion from each sample was collected as a premix to run on a mass spectrometer to ensure complete labeling. The samples were frozen at -80 °C until the results showed good labeling. At that point, the frozen samples were thawed, and the reaction was quenched by adding 3 μL of 5% hydroxylamine to each sample with incubation for 15 min at room temperature, shaking at 400 rpm. The samples within each set were combined and completely dried. Each of the 20 sets were desalted using the Discovery C18 50 mg - 1 mL solid phase extraction cartridges as described above and once again assayed to determine the final peptide concentration. An equal mass of peptide from each set was fractionated using high pH reversed-phase liquid chromatography (LC) into 24 fractions each and stored at -20 °C until LC-MS/MS analysis.

### Proteomics Analysis

A Waters nano-Acquity dual pumping ultra high-performance liquid chromatography (UPLC) system (Milford, MA) was configured for on-line trapping of a 5 µL injection at 5 µL/min for 10 min followed by reversed-flow elution onto the analytical column at 200 nL/min. The trapping column was slurry packed in-house using 360 µm o.d. x 150 um id fused silica (Polymicro Technologies Inc., Phoenix, AZ) Jupiter 5 µm C18 media (Phenomenex, Torrance, CA) with 2 mm sol-gel frits on either end. The analytical column was slurry packed in-house using Waters BEH 1.7 µm particles packed into a 35 cm long, 360 µm O.D. x 75 µm I.D. column with an integrated emitter (New Objective, Inc., Littleton, MA). Mobile phases consisted of (A) 0.1% formic acid in water and (B) 0.1% formic acid in acetonitrile with the following gradient profile (min, %B): 0, 1; 10, 8; 105, 25; 115, 35; 120, 75; 123, 95; 129, 95; 130, 50; 132, 95; 138, 95; 140,1. MS analysis was performed using a Thermo Eclipse mass spectrometer (Thermo Scientific, San Jose, CA). The ion transfer tube temperature and spray voltage were 300 °C and 2.4 kV, respectively. Data were collected for 120 min following a 27 min delay from when the trapping column was switched in line with the analytical column. MS spectra were acquired from 300-1800 m/z at a resolution of 120k (AGC target 4e5) and while the top 12 FT-HCD-MS/MS spectra were acquired in data dependent mode with an isolation window of 0.7 m/z and at an orbitrap resolution of 50K (AGC target 5e4) using a fixed collision energy (HCD) of 32 and a 30 sec exclusion time.

Tandem mass spectra were extracted and had mass errors corrected with mzRefinery [29]. Peptides were identified by searching against the SARS-CoV2 (downloaded on March 2, 2022) and human Swiss-Prot proteins (downloaded on June, 20, 2021) from the Uniprot Knowledgebase using MS-GF+ [30]. The searching parameters included 20 ppm parent mass tolerance, trypsin digestion in at least one of the peptide termini, methionine oxidation as variable modification, and cysteine carbamidomethylation and N-terminal/lysine TMT derivatization as invariable modifications. The intensities of the TMT reporter ions were extracted with MASIC [31] and used for quantitative analysis.

For the analysis of adenoviral proteins, the J&J vaccine was digested with trypsin and analyzed by LC-MS/MS as described above. Peptide sequences were determined by *de novo* sequencing using the PEAKS software (Bioinformatics Solutions Inc.). Identified peptides were blasted against the Uniprot Knowledgebase and the matched full protein sequences were retrieved and appended to the searched sequences for MS-GF+ as described above.

### Lipidomics Analysis

Total lipid extracts were analyzed using liquid chromatography electrospray ionization tandem mass spectrometry comprised of a Thermo Vanquish Flex UPLC system interfaced with a Thermo Lumos mass spectrometer. Prior to analysis, samples were reconstituted in 10% chloroform and 90% methanol, 10 µL of which was injected onto a reversed phase Waters CSH column (3.0 mm x 150 mm x 1.7 µm particle size), maintained at 42 °C and separated over a 34 min gradient (mobile phase A: acetonitrile:water (40:60) containing 10 mM ammonium acetate; mobile phase B: acetonitrile:isopropanol (10:90) containing 10 mM ammonium acetate) at a flow rate of 250 µL/min with the following gradient profile (min, %B) 0, 40; 2, 50; 3, 60; 12, 70; 15, 75; 17, 78; 19, 85; 22, 99; 34, 99. Waters Acquity UPLC CSH C18 VanGuard Pre-column (130Å, 1.7 µM packing and dimensions of 2.1 mm by 5 mm) was added to the analytical column to guard against sample particulate. Samples were analyzed in both positive and negative ionization modes using HCD (higher-energy collision dissociation) and CID (collision-induced dissociation) fragmentation mechanisms. The Fusion Lumos HESI source parameters were set as follows: spray voltage 3.5 or 3.4 kV for positive and negative modes respectively; capillary temperature 350 °C; S lens RF level 30 arbitrary units; aux gas heater temperature 350 °C; sheath, auxiliary, and sweep gas flows of 50, 10, and 1, respectively. Full MS scan data were acquired at a resolving power of 120,000 FWHM at m/z 200 with the scanning range of m/z 200–1800. The data dependent acquisition parameters used to obtain product ion spectra were as follows: loop count 12 alternating between CID and HCD, isolation width of 2 m/z units, default charge state of 1, activation Q value of 0.18 for CID, HCD resolving power of 7,500 FWHM at m/z 200, normalized collision energies for CID of 38 with detection in the ion trap and stepped collision energy of 25, 30, and 35 for HCD with detection in the orbitrap. Lipid identifications and associated integrated peak area data were generated using MS-DIAL [32]. All identifications and integrated peaks were manually validated and exported for statistical analysis.

For the detection of vaccine components in the milk samples, synthetic lipid structures and their possible tandem mass fragments were drawn in ChemSketch (ACDLabs). Tandem mass spectra and extracted-ion chromatograms were analyzed manually with Xcalibur (Thermo Scientific).

### Metabolomics Analysis

Metabolites were analyzed using reversed phase (RP) and hydrophilic interaction chromatography (HILIC) separations on a Thermo Fisher Scientific Q Exactive Plus mass spectrometer (Thermo Scientific, San Jose, CA) coupled with a Waters Acquity UPLC H class liquid chromatography system (Waters Corp., Milford, MA). Metabolite extracts were reconstituted in 100 µL of 80% LC-MS grade acetonitrile and 20% nanopure water. RP separations were performed by injecting 5 µL of sample onto a Thermo Scientific Waters Acquity UPLC BEH C18 column (130 Å, 1.7 µm, 2.1 mm ID X 100 mm L) preceded by a Acquity UPLC BEH C18 Vanguard Pre-Column (130 Å, 1.7 µm, 2.1 mm ID X 5 mm L) heated to 40 °C. Metabolites were separated using a 15-min gradient with data collected on the first 10 min. Data were acquired in both positive and negative mode with separate injections. The gradient used was identical, but the solvent composition was different between the modes. The positive mode mobile phase A consisted of 0.1% formic acid in nanopure water with the mobile phase B consisting of 0.1% formic acid in LC-MS grade methanol, while the negative mode mobile phase A consisted of 6.5 mM ammonium bicarbonate in nanopure water at a pH of 8.0 with the mobile phase B consisting of 6.5 mM ammonium bicarbonate in 95% LC-MS grade methanol and 5% nanopure water. The gradient used was as follows (min, flowrate in mL/min, %B): 0,0.35,5; 4,0.35,70; 4.5,0.35,98; 5.4,0.35,98; 5.6,0.35,0.5; 15,0.35,0.5. HILIC separations were performed by injecting 3 µL of sample onto a Waters Acquity UPLC BEH Amide column (130 Å, 1.7 µm, 2.1 mm ID X 100 mm L) preceded by a Acquity UPLC BEH Amide Vanguard Pre-Column (130 Å, 1.7 µm, 2.1 mm ID X 5 mm L) heated to 40 °C. Metabolites were separated using a 21-min gradient with data collected for the first 16 min. For HILIC, data were acquired in both positive and negative mode in separate injections using the same mobile phase compositions. The HILIC mobile phase A used consisted of 0.125% formic acid and 10 mM ammonium formate in nanopure water with a mobile phase B consisting of 0.125% formic acid and 10mM ammonium formate in 95% LC-MS grade acetonitrile and 5% nanopure water. The gradient used was as follows (min, flowrate in mL/min, %B): 0,0.4,100; 2,0.4,100; 5,0.4,70; 5.7,0.4,70; 7,0.4,40; 7.5,0.4,40; 8.25,0.4,30; 10.75,0.4,100. For both RP and HILIC separations the Thermo Fisher Scientific Q Exactive was equipped with a HESI source and high flow needle with the following parameters: spray voltage of 3.6 kV in positive mode and 3 kV in negative mode, capillary temperature of 350 °C in positive mode and 275 °C in negative mode, aux gas heater temp of 325 °C in positive mode and 300 °C in negative mode, sheath gas at 45 L/min in positive mode and 30 L/min in negative mode, auxiliary gas at 15 L/min in positive mode and 25 L/min in negative mode, and spare gas at 1L/min in positive mode and 2 L/min in negative mode. Metabolites were analyzed at a resolution of 70 k and a scan range of 70 to 1000 m/z for parent ions followed by data dependent MS/MS HCD fragmentation on the top 4 ions with a resolution of 17.5 K and stepped normalized collision energies of 20, 30, and 40. Metabolite identifications and associated integrated peak area data were generated using MS-DIAL. All identifications and integrated peaks were manually validated and exported for statistical analysis.

### Statistical and Pathway Analyses

Statistical analysis first included quality control analysis, removing any lipids, metabolites, and peptides that were not identified in enough samples for a repeated measures analysis of variance (rANOVA) model. A sample-sample correlation and principal component analysis were utilized to assess any sample level issues and one sample, a subject receiving the Moderna vaccine at the third time point, showed poor correlation with the sample set. Further evaluation identified a data processing issue where the lipids software stopped collecting data during the analysis. This was the only sample removed prior to statistical analysis from the lipid data. There were no identified outliers in the metabolomics data. The positive and negative mode lipidomics data, each metabolomic dataset, and the proteomics data were then normalized to median abundance.

To evaluate the change in milk composition over time between the vaccinated women rANOVA was performed where lipid, metabolite or protein abundance is the response variable, time is the within subject variable, and the vaccine is the predictor variable. For the comparison of the milk produced by women with COVID-19 versus that produced by vaccinated women a two-sample t-test was utilized. Given that many lipids, metabolites, and proteins are highly correlated a meta-analysis approach to multiple test correction was applied [33] using R package *poolr* by Cinar and Viechtbauer (2022), estimating the correction using the correlation matrix for each pair of biomolecules. Finally, an adjusted p-value threshold that controls the family-wise error rate to 0.05 using the Sidak multiple comparison. The functional pathway enrichment analysis was done using DAVID Knowledgebase [34]. Only the KEGG pathways were used for our analysis, and they were filtered with an enrichment p-value ≤ 0.05.

**Complete proteomics, lipidomics, and metabolomics data can be found at:** https://figshare.com/account/home#/projects/200344

---

**Supplemental Figure 1.**
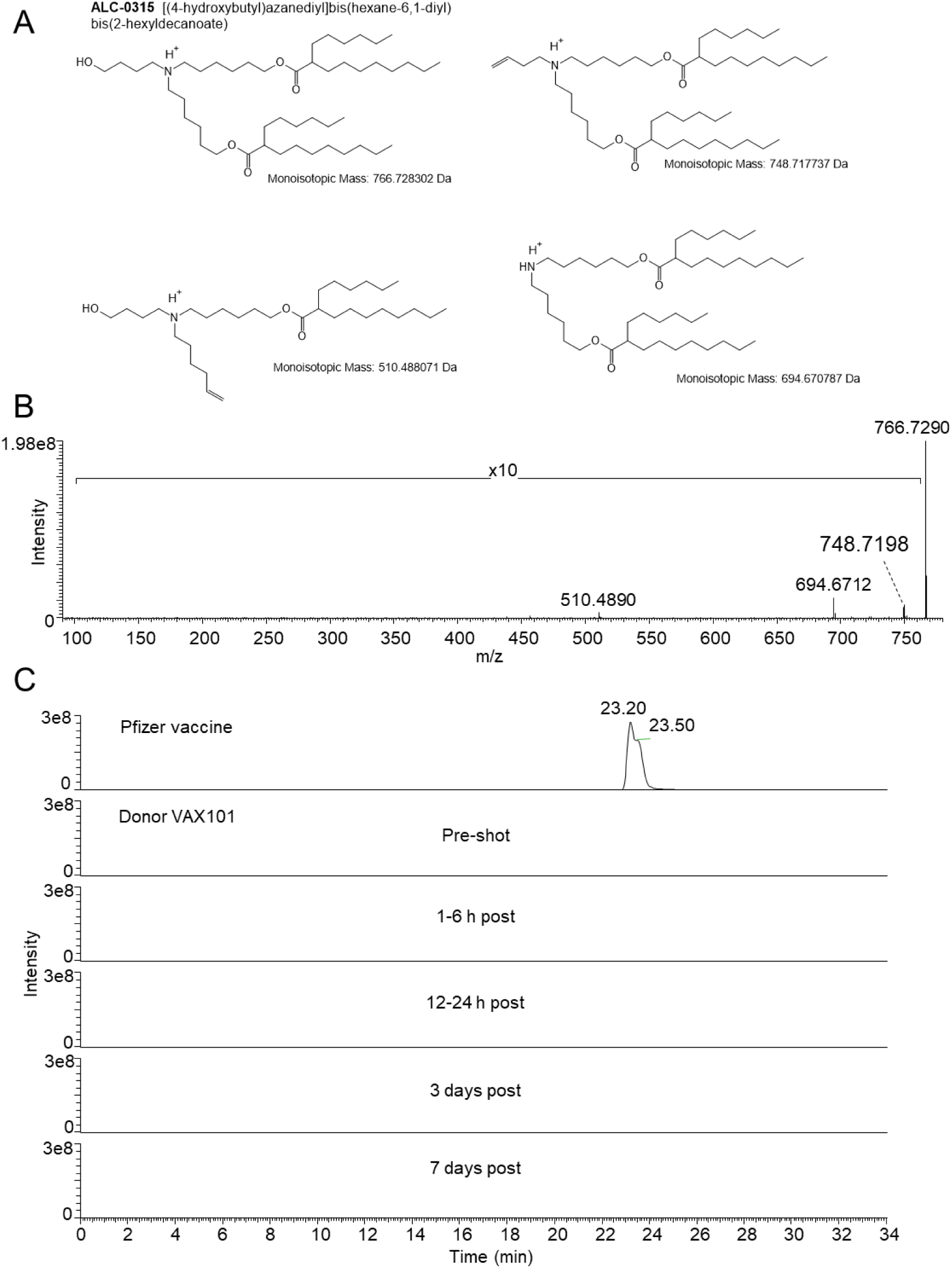
Analysis of the Pfizer vaccine component ALC-0315. (A) Structure of ALC-0315 and predicted fragments that can be generated in the mass spectrometer. (B) Tandem mass spectrum of ALC-0315. (C) Extracted-ion chromatogram of ALC-0315 in the data from donor VAX101. Importantly, the peak ALC-0135 was not detected in any of the milk samples when analyzed with MS Dial.

**Supplemental Figure 2.**
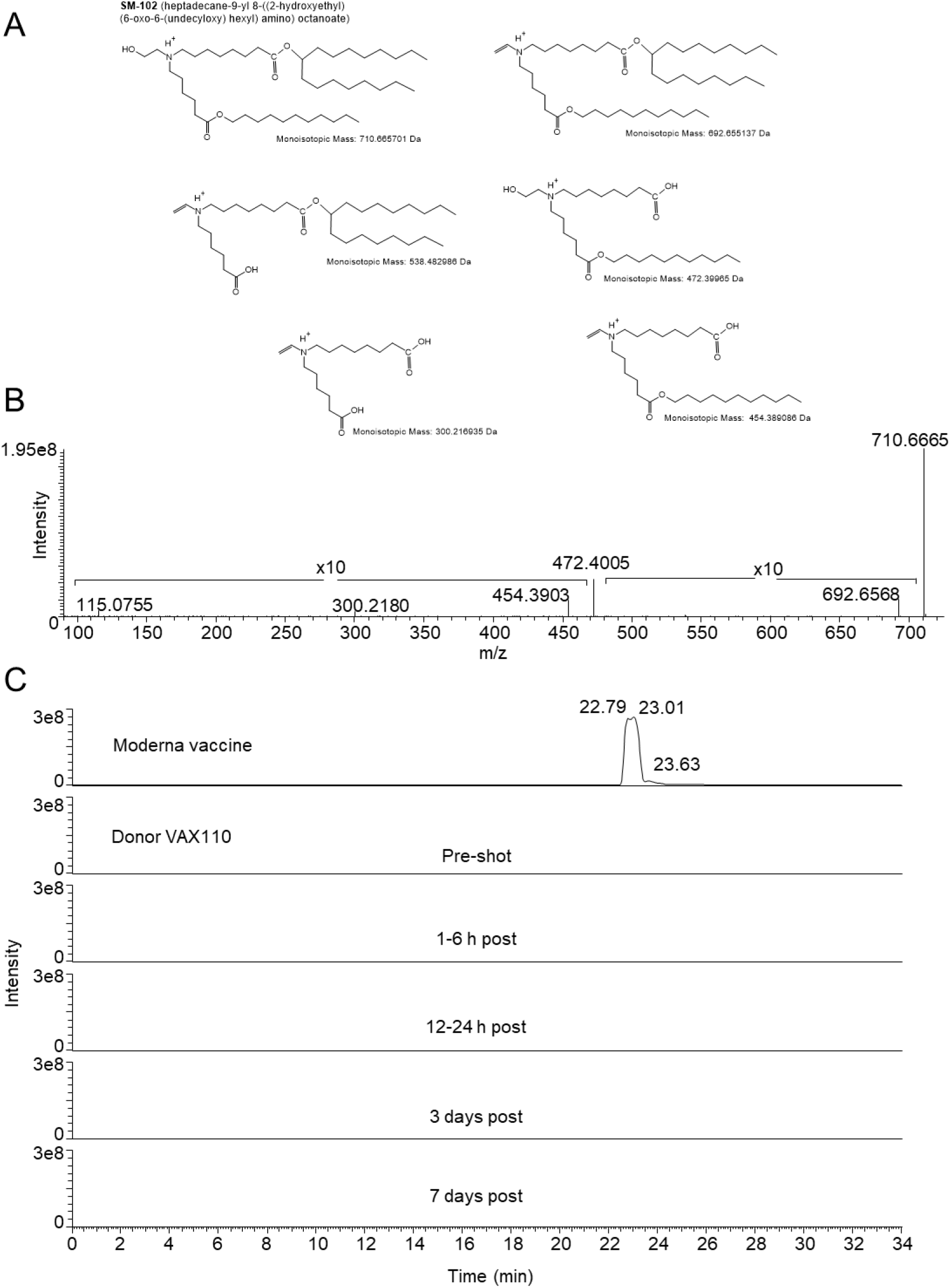
Analysis of the Moderna vaccine component SM-102. (A) Structure of SM-102 and predicted fragments that can be generated in the mass spectrometer. (B) Tandem mass spectrum of SM-102. (C) Extracted-ion chromatogram of SM-102 in the data from donor VAX110. Importantly, the peak SM-102 was not detected in any of the milk samples when analyzed with MS Dial.

**Supplemental Table 1:** Type of vaccine administered, age, months postpartum, and race/ethnicity for vaccinated participants and days since COVID-19 diagnosis, age, months postpartum, and race/ethnicity for infected participants.

| Vaccinated Participant ID | Vaccine | Age | Months postpartum | Race/Ethnicity |
| --- | --- | --- | --- | --- |
| VAX101a | Pfizer | 36 | 2 | White, Non-Hispanic |
| VAX107a | Pfizer | 33 | 7 | White, Non-Hispanic |
| VAX108a | Pfizer | 34 | 11 | Asian or Pacific Islander |
| VAX109a | Pfizer | 41 | 8 | White, Non-Hispanic |
| VAX119a | Pfizer | 37 | 11 | White, Non-Hispanic |
| VAX136a | Pfizer | NA | NA | NA |
| VAX144a | Pfizer | 31 | 7 | White, Non-Hispanic |
| VAX166a | Pfizer | 47 | 7 | White, Non-Hispanic |
| VAX180a | Pfizer | 32 | 1 | White, Non-Hispanic |
| VAX182b | Pfizer | 34 | 9 | White, Non-Hispanic |
| VAX186a | Pfizer | NA | < 1 | NA |
| VAX187a | Pfizer | 35 | 2 | Asian or Pacific Islander |
| VAX191a | Pfizer | 31 | 4 | Asian or Pacific Islander |
| VAX195a | Pfizer | 36 | 6 | Asian or Pacific Islander |
| VAX205a | Pfizer | 32 | 8 | Asian or Pacific Islander |
| VAX206b | Pfizer | 38 | 11 | White, Non-Hispanic |
| VAX207a | Pfizer | 39 | 21 | White, Non-Hispanic |
| VAX208a | Pfizer | 35 | 3 | White, Non-Hispanic |
| VAX210a | Pfizer | 34 | 10 | White, Non-Hispanic |
| VAX212a | Pfizer | 27 | 5 | White, Non-Hispanic |
| VAX213a | Pfizer | 35 | 4 | White, Non-Hispanic |
| VAX217a | Pfizer | 31 | 12 | White, Non-Hispanic |
| VAX221a | Pfizer | 33 | 1 | White, Non-Hispanic |
| VAX226a | Pfizer | 32 | < 1 | White, Non-Hispanic |
| VAX237a | Pfizer | 32 | 5 | White, Non-Hispanic |
| VAX110a | Moderna | 37 | < 1 | White, Non-Hispanic |
| VAX112c | Moderna | 37 | 6 | White, Non-Hispanic |
| VAX114a | Moderna | 30 | 12 | Multiracial |
| VAX117a | Moderna | 34 | 5 | White, Non-Hispanic |
| VAX133a | Moderna | 34 | 7 | White, Non-Hispanic |
| VAX134a | Moderna | NA | < 1 | NA |
| VAX150a | Moderna | 38 | 2 | White, Non-Hispanic |
| VAX169a | Moderna | 31 | 5 | White, Non-Hispanic |
| VAX185a | Moderna | 38 | 11 | White, Non-Hispanic |
| VAX190a | Moderna | 39 | 2 | Multiracial |
| VAX192a | Moderna | 34 | 4 | White, Non-Hispanic |
| VAX198a | Moderna | 33 | 2 | Asian or Pacific Islander |
| VAX211a | Moderna | 33 | < 1 | White, Non-Hispanic |
| VAX214a | Moderna | 34 | 35 | White, Non-Hispanic |

| VAX216a | Moderna | 37 | 4 | White, Non-Hispanic |
| --- | --- | --- | --- | --- |
| VAX227a | Moderna | 36 | 18 | White, Non-Hispanic |
| VAX228a | Moderna | NA | NA | NA |
| VAX231a | Moderna | 33 | 12 | White, Non-Hispanic |
| VAX232a | Moderna | 32 | 9 | White, Non-Hispanic |
| VAX233a | Moderna | 33 | 10 | White, Non-Hispanic |
| VAX234a | Moderna | 39 | 9 | White, Non-Hispanic |
| VAX235a | Moderna | 37 | 8 | White, Non-Hispanic |
| VAX236a | Moderna | 40 | 1 | White, Non-Hispanic |
| VAX148a-J | J&J | 38 | 3 | White, Non-Hispanic |
| VAX149a-J | J&J | 36 | 7 | White, Non-Hispanic |
| VAX151a-J | J&J | 33 | 8 | White, Non-Hispanic |
| VAX153a -J | J&J | 30 | 6 | White, Non-Hispanic |
| VAX154a-J | J&J | 41 | 3 | White, Non-Hispanic |
| VAX163a-J | J&J | 35 | < 1 | White, Non-Hispanic |
| VAX164a-J | J&J | 32 | 13 | Asian or Pacific Islander |
| VAX165a-J | J&J | 28 | 17 | Multiracial |
| VAX171a-J | J&J | 32 | 8 | White, Non-Hispanic |
| VAX172a-J | J&J | NA | NA | White, Hispanic or Latino |
| VAX173a-J | J&J | 31 | 7 | White, Non-Hispanic |
| VAX177a-J | J&J | 30 | 7 | White, Non-Hispanic |
| COV406d | J&J | 34 | 14 | White, Hispanic or Latino |
| **Infected Participant ID** | **Days post positive test** | **Age** | **Months postpartum** | **Race/Ethnicity** |
| LC1 | 2 | 40 | 8 | White, Non-Hispanic |
| LC21 | 2 | 26 | 5 | White, Non-Hispanic |
| LC23 | 3 | 26 | 6 | White, Non-Hispanic |
| LC25 | 6 | 41 | 17 | White, Non-Hispanic |
| LC26 | 12 | 30 | < 1 | White, Non-Hispanic |
| LC29 | 3 | 37 | 30 | White, Hispanic or Latino |
| LC31 | 1 | 39 | 10 | White, Non-Hispanic |
| LC32 | 2 | 32 | 4 | White, Non-Hispanic |
| LC34 | 3 | 36 | 4 | White, Non-Hispanic |
| LC37 | 9 | 38 | 2 | White, Non-Hispanic |
| LC4 | 4 | 32 | 8 | White, Non-Hispanic |
| LC40 | 4 | 32 | 13 | White, Non-Hispanic |
| LC41 | 3 | 31 | < 1 | White, Non-Hispanic |
| LC42 | 3 | 39 | 7 | White, Non-Hispanic |
| LC44 | 4 | 33 | 5 | White, Non-Hispanic |
| LC45 | 5 | 36 | 7 | White, Non-Hispanic |
| LC47 | 6 | 26 | 13 | White, Non-Hispanic |
| LC48 | 5 | 26 | 6 | White, Hispanic or Latino |
| LC50 | 8 | 30 | 2 | White, Non-Hispanic |
| LC52 | 8 | 30 | < 1 | White, Non-Hispanic |
| LC53 | 5 | 33 | 3 | White, Hispanic or Latino |
| LC54 | 6 | 29 | 8 | White, Non-Hispanic |
| LC55 | 5 | 36 | 21 | White, Non-Hispanic |
| LC6 | 3 | 29 | 3 | White, Non-Hispanic |
| LC8 | 5 | 30 | 7 | White, Hispanic or Latino |

